# S2′ cleavage site plays a decisive role in expansion of Vero cell tropism by infectious bronchitis virus HV80, with Q855H promoting cell-to-cell fusion

**DOI:** 10.1101/2023.04.19.537537

**Authors:** Yi Jiang, Xu Cheng, Mingyan Gao, Xinhong Dou, Yan Yu, Haiyu Shen, Mengjun Tang, Sheng Zhou, Daxin Peng

## Abstract

Infectious bronchitis virus (IBV) has restricted cell tropism. Apart from the Beaudette strain, other IBVs cannot infect mammalian cell lines. The limited cell tropism of other IBVs has hindered the development of IBV vaccines and research on mechanisms of IBV infection. In a previous study, a new Vero-cell-adapted strain HV80 was obtained via serial chicken embryo and cell passaging of strain H120 and 17 mutations leading to amino acid substitutions occurred in replication gene 1a, S gene and E gene. This study, we constructed recombinants that expressed chimeric S glycoprotein, S1 or S2 subunit of strain H120, and demonstrated that mutations in S2 subunit were related to the Vero cell adaption of strain HV80. With a genome backbone of strain HV80 or H120, and expression of chimeric S2′ cleavage site of H120 or HV80, two recombinants demonstrated that the RRRR_690_/S motif at the S2′ cleavage site played a key role in Vero cell adaption of strain HV80. Another six amino acid substitutions in the S2 subunit of the recombinants showed that F692V enhanced the capability of invasion of HV80 strain, and Q855H induced the formation of syncytia. A transient transfection assay demonstrated different mechanisms for virus-to-cell fusion and cell-to-cell fusion induced by S glycoprotein. The PRRR_690_/S motif at the S2′ cleavage site could be activated by proteases in the process of cell-to-cell fusion, while H855Q substitution did not affect the cell invasion of HV80, but hindered the cell-to-cell fusion by blocking activation of the S2′ cleavage site.

**IMPORTANCE:** Infectious bronchitis is an acute respiratory disease that has caused large economic losses to the poultry industry. As a member of the gamma-coronaviruses, the restricted cell tropism of infectious bronchitis virus (IBV) limits the development of cellular vaccines and research on infection mechanisms. As a strain that can replicate effectively in mammalian cell lines, studies of HV80’s adaptive mechanisms point a way for engineering other IBVs for adaptation in mammalian cell lines. In our study, different recombinants were constructed by reverse genetic techniques, and demonstrated the different mechanism between virus-to-cell and cell-to-cell fusion induced by HV80 S glycoprotein. The acquisition of a highly efficient S2′ cleavage site enabled the virus to invade Vero cells. The Q855H substitution played a key role in cell-to-cell fusion, and provided a more efficient model of infection in Vero cells. Our study provides new theoretical insights into mechanisms of IBV adaptation in mammalian cell lines.

---

Infectious bronchitis virus (IBV) belongs to the order Nidovirales, family Coronaviriae, sub-family Coronavirinae within the genus gamma-coronavirus. It is a positive-strand RNA virus with a genome of 27.6 kb, which consists of structural proteins such as spike (S), membrane (M), envelope (E) and nucleocapsid (N) (1). Coronaviruses (CoVs) have a wide range of hosts and tissue tropism, and infect birds and mammals, including humans, pigs, dogs, cats, bats and chickens (2). This host range and tissue tropism are associated with the mechanism of viral invasion of host cells, which is determined by S glycoprotein (3).

In the process of viral infection, S glycoprotein is cleaved into two subunits, S1 and S2, which respectively participate in two key events: (1) S1 subunit binds to the receptor on the cell membrane and (2) S2 subunit induces virus-to-cell and cell-to-cell fusion (4). S1 and S2 subunits exist on the virion surface as a trimer. The receptor binding domain (RBD) is located in the S1 subunit, and can be further subdivided into the N-terminal domain (NTD) and C-terminal domain (CTD), according to the structure and function (5). The location of RBD for CoVs differs; the CTD of alpha- and beta-CoVs, such as human coronavirus 229E (HCoV-299E), transmissible gastroenteritis virus (TGEV), feline coronaviruses (FCoV), SARS-CoV, MERS-CoV and SARS-CoV-2, binds to protein receptors, including peptidase N, angiotensin-converting enzyme 2 and dipeptidyl peptidase 4 (6–12). Except for carcinoembryonic antigen-related cell adhesion molecule 1 (CEACAM1) protein receptor, NTD appears to mainly bind glycans, including sialic acids and O-acetylated sialic acids (13–16). For gamma-CoVs such as IBV, the protein receptor is still unknown, and the NTD of IBV S glycoprotein can bind to sialic acid, which plays an important role during virus entry (17, 18). Unlike other IBVs, the Beaudette strain is a special strain that has ability to infect some mammalian cell lines, such as Vero and BHK-21 (19, 20). Some studies indicated that expansion of cell tropism of the Beaudette strain was not related to S1 subunit. Binding to heparan sulfate (HS) may be one reason for adaption of the Beaudette strain to mammalian cell lines, and the binding domain is located at the S2′ cleavage site (21).

After receptor engagement, IBV needs to induce fusion of viral and cellular membranes by proteolytic action of cellular protease. At present, two protease cleavage sites, S1/S2 and S2′ have been identified in S glycoprotein. The S1/S2 cleavage site is located at the boundary between the S1 and S2 subunits, while S2′the cleavage site is located immediately upstream of the fusion peptide (1). The S2′ cleavage site of SARS-CoV, MERS-CoV and SARS-CoV-2 can be proteolytically cleaved by transmembrane serine protease 2 at the plasma membrane (12, 22, 23). Unlike SARS-CoV, a polybasic residue motif is inserted at the boundary between the S1 and S2 subunits. This S1/S2 cleavage site render SARS-CoV-2 prone to cleavage by furin during viral biogenesis at the endoplasmic reticulum/Golgi and trans-Golgi compartments (24, 25). Two furin cleavage site (S1/S2 and S2’) located on S glycoprotein of MERS-CoV, the S1/S2 boundary site is cleaved by furin during biosynthesis in the host cell, and the S2′ site cleavage occurs during virus infection(26). Most IBVs only have the S1/S2 furin cleavage site, expect for the Beaudette strain, which has the second furin cleavage site at S2′. Unlike other IBVs, activation of the S2′ cleavage site may be the key for the Beaudette strain to extend its tropism for some mammalian cells (27, 28). Furin cleavage site insertion into IBV YN strain demonstrated that S2′ cleavage site plays a key role in progression of central nervous system damage (29, 30).

As a class I viral fusion protein, after activation of host proteolytic protease, the fusion peptide (FP), two heptad repeat regions (HR1 and HR2) from S2, plays a role in membrane fusion (Figure 1). A second proteolytic cleavage site on S2′ triggers conformational changes to expose the FP as a cell membrane target (31). FP is composed of a short segment, conserved across the viral family, which is highly hydrophobic (32, 33). One amino acid substitution of L803A, L804A or F805A on FP of SARS-CoV prevents fusion (34). HR1 is downstream of the FP, while HR2 is upstream of the transmembrane domain, and both are composed of repetitive heptapeptides (35). After insertion of the FP, the trimeric HR2 region folds back to the hydrophobic grooves of HR1 trimer, and forms a six-helix bundle. The six-helix bundle damages the stability of the lipid bilayers, and results in conformational rearrangement, which pulls the virus and host cell membrane into proximity to form a fusion pore. The viral genome is released into the cytoplasm through this fusion pore (36–38). Mutation of M936V, P939L, F948L and S949I appeared on the S glycoprotein of a serially passaged MHV-59 strain. These amino acids in or adjacent to the HR1 region led to expansion of the host range of the V51 variant to other mammalian cell lines *in vitro* (39, 40). Almost IBV strains can only replicated in primary cells, such as CECK and CK, thus the pathogenic mechanism studies at cellular level focused on the Beaudette strain, a Vero-cell-adapted strain. The Beaudette strain was adapted to Vero cells by serial propagation in embryonated eggs and then Vero cells for 65 passages. Forty-nine mutations appeared on the genome of p65, and 26 substitutions were located in the S glycoprotein (41). Among them, F857L substitution in HR1 domain of Vero-cell-adapted strain p65 lost the ability for cell membrane fusion (42).

**FIG 1.**
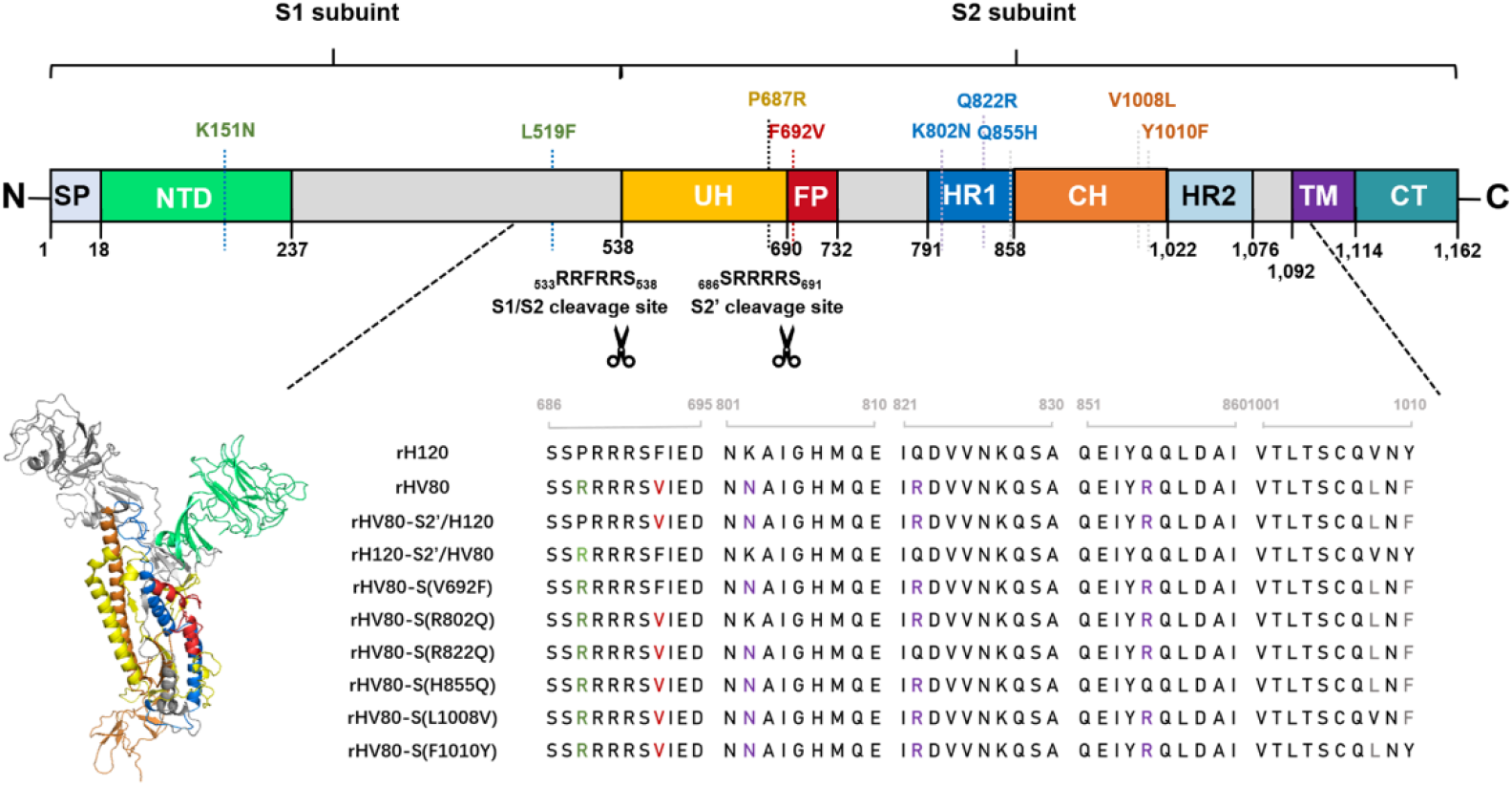
Summary of amino acid substitutions in the IBV S genes from Vero-cell-adapted HV80 strains. (A) Schematic diagram of the IBV HV80 strain (GenBank accession number: OP684009) S gene structure and functional domains. SP, signal peptide; NTD, N-terminal domain; S1/S2, S1/S2 cleavage site; UH, upstream helix; S2′, S2′ cleavage site; FP, putative fusion peptide; HR1, heptad repeat 1; HR2, heptad repeat 2; TM, transmembrane domain; and CT, C-terminal domain. Black dotted lines indicate nine amino acid substitutions occurred on S gene of HV80 strain compared with its parental strain H120. (B) The structure prediction of the monomer of HV80 spike ectodomain in the prefusion conformation. Cryoelectron microscopy structure in the prefusion conformation was predicted by the Swiss-model referred to 6cv0.1. The different components of S glycoprotein are colored differently, which is consistent with the schematic illustration (A). (C) The partial amino acid sequence of the S2 subunit showed from different substitution recombinants.

In a previous study(43), a new Vero-cell-adapted strain, HV80, was obtained via serial passage in chicken embryo and primary CK and Vero cells of strain H120. Whole genome sequencing of strain HV80 showed that 17 mutations leading to amino acid substitutions appeared in the replicase 1a, S and E gene. Of these, 9 mutations occurred in the S gene. We constructed a series of IBV recombinants under consideration of these amino acid substitutions and the domains in which they reside. Indirect immunofluorescence assay and viral growth curves in infected Vero cells showed that the acquisition of a highly efficient S2’ cleavage site enabled the virus to invade Vero cells, and the Q855H substitution played a key role in cell-to-cell fusion, and provided a more efficient model of infection in Vero cells. A transient transfection assay demonstrated the PRRR_690_/S motif at the S2′ cleavage site could be activated by proteases in the process of cell-to-cell fusion. H855Q substitution did not affect the cell invasiveness of HV80, but hindered cell-to-cell fusion by blocking activation of the S2′ cleavage site. This study extends our understanding of the adaptive mechanism of IBV in mammalian cell lines, and indicates the possibility of engineering other IBV strains for adaptation in mammalian cell lines.

## RESULTS

### S glycoprotein determined the expansion of cell tropism in Vero cells, and the S2 subunit played an important role in adaption of strain HV80 to Vero cells

IFA showed that green fluorescence was not visible in Vero cells infected with rH120 strain at 36 h post-infection (hpi). In contrast, a large amount of green fluorescence with large areas of cell fusion appeared in Vero cells infected with rHV80 strain (Fig. 2A). The viral growth curves showed that viral RNA content from the supernatant of Vero cells infected with rHV80 increased gradually over time, and reached a peak at 60 hpi. No increase in viral copies from the supernatant indicated that strain rH120 could not infect Vero cells (Fig. 2B and C). Within the genomic background derived from strain HV80, the recombinant strain rHV80-S/H120, expressing chimeric S glycoprotein of H120, was constructed using a reverse genetic system. Green fluorescence was not detected in Vero cells infected with strain rHV80-S/H120, and the virus could not efficiently replicate within Vero cells, which indicated that the S gene had been replaced with the corresponding gene from H120 and lost the ability to infect Vero cells (Fig. 2A and B). We constructed rHV80-S1/H120 and rHV80-S2/H120, which expressed the chimeric S1 or S2 subunit of the H120 strain, with the genome backbone of strain HV80. At 36 hpi, a large amount of green fluorescence appeared in some single cells or syncytia of Vero cells infected with rHV80-S1/H120. In contrast, there was no green fluorescence in Vero cells infected with rHV80-S2/H120. Growth curves showed that rHV80-S2/H120 did not replicate in Vero cells, the ability of rHV80-S1/H120 strain to infect and replicate in Vero cells was not affected by the S1 subunit replacement (Fig. 2A and C).

**FIG 2.**
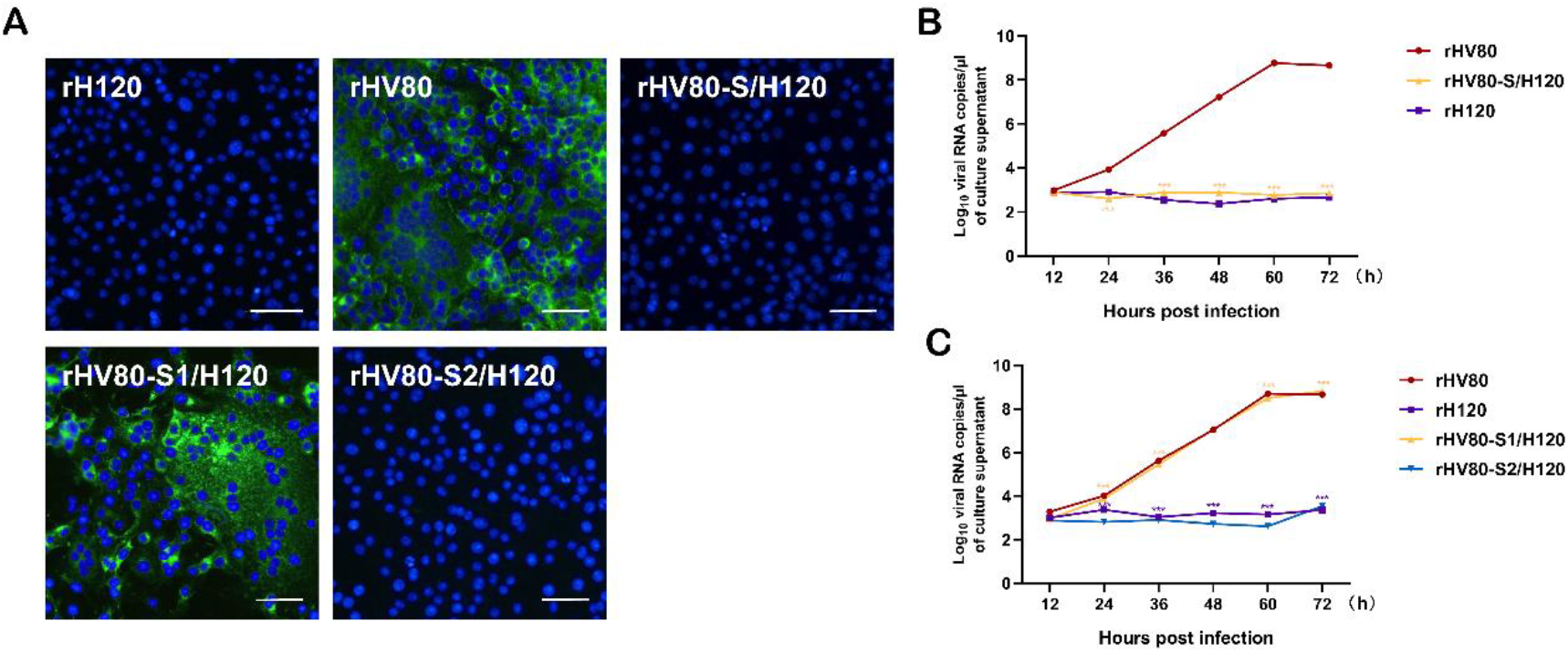
Vero cells infected with different recombinant chimeric viruses expressing the S glycoprotein or S1/S2 subunit. (A) Vero cells were grown in six-well plates for 24 h and infected with rH120, rHV80, rHV80-S/H120, rHV80-S1/H120 or rHV80-S2/H120 at 10^7^copies/100 μl. After 24 h of infection, the cells were fixed with cold methanol for IFA assay. Nuclei were labeled with DAPI (blue). Bar, 50 μm. (B) and (C) Growth curves for different recombinant viruses in Vero cells. Vero cells in 12-well plates were inoculated with different recombinant viruses. The supernatant was harvested at 12, 24, 36, 48, 60 and 72 hpi. Viral RNA copies were quantified by real-time RT-PCR. Error bars indicate the standard deviation.

### In the S glycoprotein of strain HV80, the S2′ cleavage site played a crucial role in adaptation to Vero cells, but rH120-S2′/HV80 showed limited capacity for infection

There was no green fluorescence in Vero cells infected with strain rHV80-S2′/H120 at 36 hpi. Only an extremely small amount of green fluorescence appeared in Vero cells infected with strain rH120-S2′/HV80 (Fig. 3A). The viral growth curves showed that there was no significant increase in viral copies from the supernatant of Vero cells infected with both the rHV80-S2′/H120 and rH120-S2′/HV80 strains, which was similar to cells infected with the rH120 strain (Fig. 3B).

**FIG 3.**
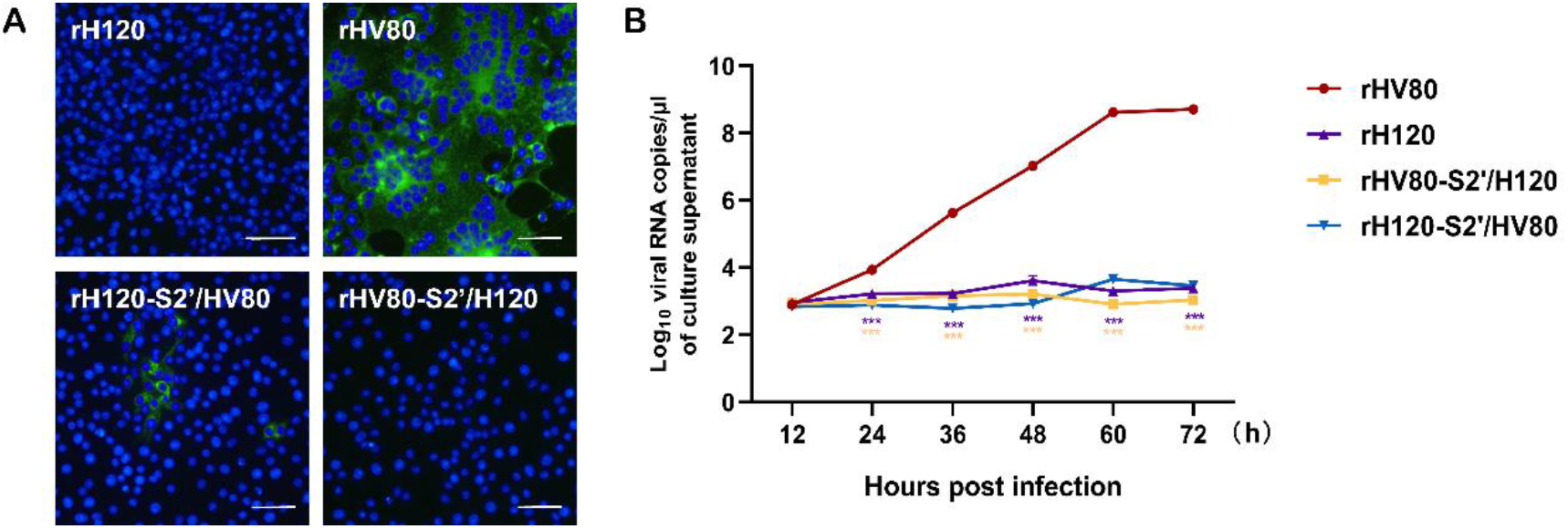
Vero cells infected with different recombinant viruses associated with the S2′ cleavage site. (A) Vero cells were grown in six-well plates for 24 h and infected with rH120, rHV80, rH120-S2’/HV80, or rHV80-S2′/H120 at 10^7^copies/100 μl. After 24 h of infection, the cells were fixed with cold methanol for IFA assay. Nuclei were labeled with DAPI (blue). Bar, 50 μm. (B) Growth curves for different recombinant viruses in Vero cells. Vero cells in 12-well plates were inoculated with different recombinant viruses. The supernatant was harvested at 12, 24, 36, 48, 60 and 72 hpi. Viral RNA copies were quantified by real-time RT-PCR. Error bars indicate the standard deviation.

### Almost all amino acid substitution affected viral replication in Vero cells, and substitution of H855Q in the S glycoprotein of HV80 was different from other amino acid substitutions in terms of green fluorescence distribution and virus growth tendency

Variable degrees of green fluorescence were observed in the Vero cells infected with six recombinants with different one-point amino acid substitutions, but the range of fluorescence was lower than that in the parental strain rHV80. Compared to other one-point substitution recombinants, a greater extent of green fluorescence was observed in Vero cells infected with rHV80-S(N802K), rHV80-S(L1008V), and rHV80-S(F1010Y) strains. Green fluorescence with a small number of moderate-sized syncytia appeared in Vero cells infected with the rHV80-S(R822Q) and rHV80-S(V692F) strains. The green fluorescence distributed as single cells were appeared in rHV80-S(H855Q) group, and no cell-to-cell fusion was observed (Fig. 4A). In viral growth curves, the viral RNA copies in the supernatant of rHV80-infected cells peaked 12 hours earlier than that of the previous growth curves due to the 10-fold increase in inoculum dose. Compared with their parental strain rHV80, the replication efficiency of other one-point substitution recombinants significantly decreased, and the time to reach replication peak was later than that in rHV80-infected cells. Except for the rHV80-S(H855Q) group, the general trends in growth curves were similar in the other five one-amino-acid substitution groups, but the speed of replication differed. Replication efficiency was similar in cells infected with rHV80-S(N802K) or rHV80-S(F1010Y), which showed faster growth than in cells infected with other substitution recombinants. Cells infected with rHV80-S(V692F) had the slowest growth. Unlike other one point substitution groups, the viral RNA copy number in the cell culture supernatant from the rHV80-S(H855Q) group was higher during the early phase of infection (24 and 36 hpi), but the peak number of viral RNA copies at 48 and 60 hpi was lower than for the other recombinant-infected groups. Strain rH120 still had no ability to replicate in Vero cells (Fig. 4B).

**FIG 4.**
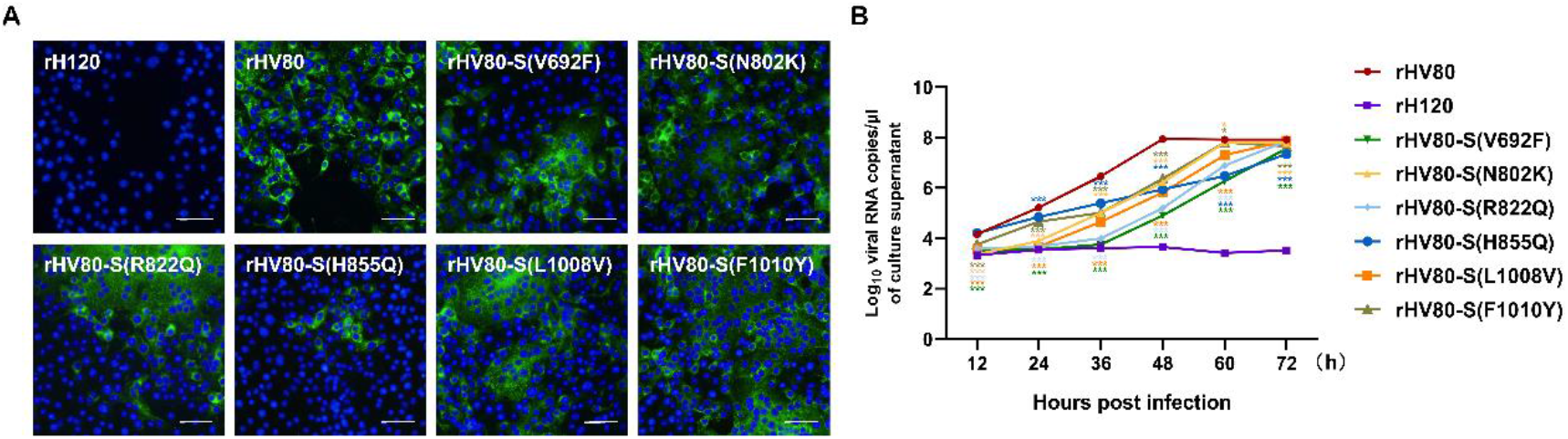
Vero cells infected with one-amino-acid substitution viruses. (A) Vero cells were grown in six-well plates for 24 h and infected with rH120, rHV80, rHV80-S2(V692F), rHV80-S2(N802K), rHV80-S2(R822Q), rHV80-S2(H855Q), rHV80-S2(L1008V) or rHV80-S2(F1010Y) at 10^8^copies/100 μl. After 24 h of infection, the cells were fixed with cold methanol for IFA assay. Nuclei were labeled with DAPI (blue). Bar, 50 μm. (B) Growth curves for different recombinant viruses in Vero cells. Vero cells in 12-well plates were inoculated with different recombinant viruses at 5×10^7^ copies/50 μl. The supernatant was harvested at 12, 24, 36, 48, 60 and 72 hpi. Viral RNA copies were quantified by real-time RT-PCR. Error bars indicate the standard deviation.

### Cell to cell spread was inhibited in Vero cells infected with rHV80-S(H855Q), and no fusion occurred in Vero cells expressing S(H120-S2′/HV80), S(HV80-V692F) or S(HV80-H855Q) with EGFP tag

We performed a cell-plaque assay to compare the plaque phenotypes of cells infected with rHV80-S2′/H120, rHV80-S(H855Q), rHV80-S(V692F) or their parental strains rHV80 and rH120. Compared with the rHV80-infected group, the number of plaques decreased in the cells infected with the rHV80-S(V692F) strain, after crystal violet staining at 72 hpi. The size of plaques in Vero cells infected with rHV80-S(H855Q) was significantly smaller than that in cells infected with rHV80 (Fig. 5A and B). No viral plaques appeared in Vero cells infected with the rHV80-S(R687P) strain (rHV80-S2’/H120).

**FIG 5.**
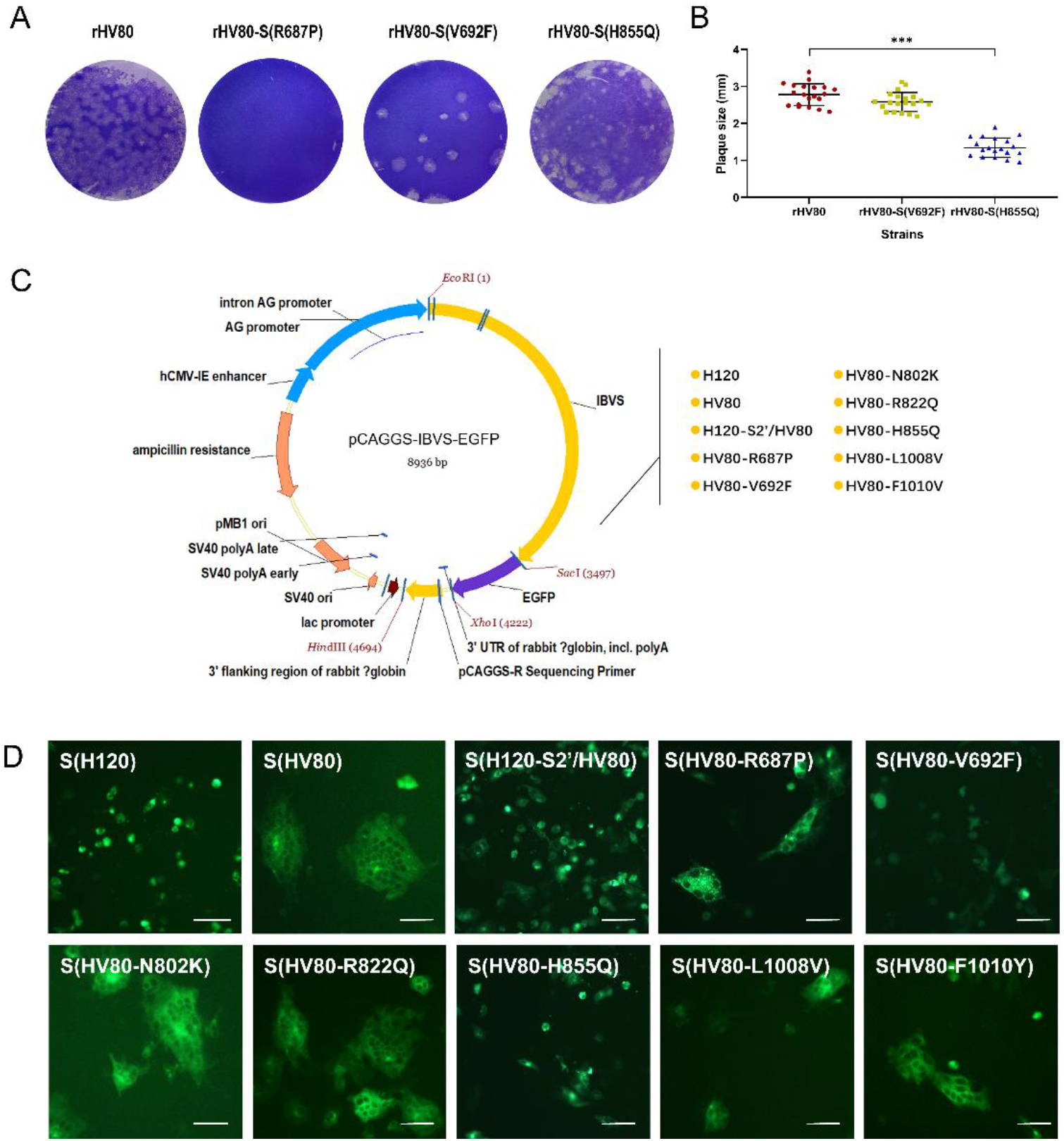
Cell-plaque assay of Vero cells infected with recombinants and cell-to-cell fusion induced by different S glycoproteins fused with EGFP-tag constructs. (A and B) Vero cells were inoculated with rHV80, rHV80-S(R687P), rHV80-S2(V692F) or rHV80-S2(H855Q). At 72 hpi, the infected cells were fixed with 4% formaldehyde solution and stained with crystal violet for 20 min. After staining, the samples were photographed and recorded (A), and the diameter of plaques was measured (B). (C) Schematic diagram of recombinant plasmid construction. Upon cleaving the vector pCAGGS-MCS with restriction enzymes (*Sac*I and *Xho*I), the EGFP fragment was inserted to construct the plasmid pCAGGS-EGFP. The target fragments of different IBV genes were inserted into the plasmid pCAGGS-EGFP using two restriction enzymes (*Eco*RI and *Sac*I). (D) Vero cells were transfected with different S glycoprotein constructs. At 36 h post-transfection, the green fluorescence and the fusion of cells transfected with pCAGGS-IBVS-EGFP plasmids were visualized under a fluorescent microscope.

The vector pCAGGS-EGFP was used as the backbone to construct the recombinant plasmids encoding the S glycoprotein from different recombinants (Fig. 5C). Vero cells were transiently transfected with the plasmid carrying S– EGFP. At 36 h after transfection, fluorescence was observed under a microscope. A large amount of green fluorescence appeared in the cells expressing H120S–EGFP, and fluorescence-positive cells were present as individual cells with high fluorescence intensity. In the cells expressing HV80S– EGFP, the fluorescence intensity was lower, and the fluorescence-positive cells were in a confluent state with syncytia of different sizes. Among the recombinant plasmid encoding S glycoprotein of the one-amino acid substitution strains, cell-to-cell fusion occurred in the positive cells expressing plasmids pCAGGS-S(HV80-N802K)-EFGP, pCAGGS-(HV80S-R822Q)-EFGP, pCAGGS-S(HV80-L1008V)-EFGP, and pCAGGS-S(HV80-F1010Y)-EFGP, which was similar to the pCAGGS-S(HV80)-EGFP transfection group. However, the fluorescence-positive cells transfected with pCAGGS-S(H120-S2′/HV80)-EFGP, pCAGGS-S(HV80-V692F)-EFGP or pCAGGS-S(HV80-H855Q)-EFGP, presented as individual cells, which was similar to the pCAGGS-S(H120)-EGFP transfection group. A small number of syncytia were present in the pCAGGS-S(HV80-R687P)-EFGP transfection group (Fig. 5D).

### Cell-to-cell fusion was significantly inhibited by the H855Q substitution, which might have been related to no activation at S2’ cleavage site

To analyze S glycoprotein cleavage, the S gene from different strains was inserted into plasmid pCAGGS-Fc (Figure 6A). Six recombinant plasmids were transfected into Vero cells. At 36 h after transfection, cell fluorescence appeared in each transfection group, which indicated S glycoprotein fusion with the Fc-tag. Almost no fusion occurred in Vero cells transfected with pCAGGS-S(H120)-Fc, pCAGGS-S(H120-S2’/HV80)-Fc, or pCAGGS-S(HV80-H855Q)-Fc. Some large syncytia were formed in Vero cells transfected with pCAGGS-S(HV80)-Fc or pCAGGS-S(HV80-V692F)-Fc. A small number of syncytia were also present in the pCAGGS-S(HV80-R687P)-Fc transfection group (Figure 6B and C). Western blotting showed that three bands of different sizes were detected in Vero cells expressing S(HV80)-Fc protein. The expression and cleavage of S(HV80)-Fc protein increased over time, and peaked at 36–48 h post transfection (Figure 6D). The cells were transfected with different recombinant plasmids, and cell lysates were harvested to analyze the cleavage of S glycoprotein at 36 h post-transfection. As in the S(HV80) transfection group, three bands of different size (S near 140 kDa, S2 near 115 kDa and S20 near 80 kDa) were detected in the S(HV80-R687P) and S(HV80-V692F) transfection groups. However, only two bands (S near 140 kDa and S2 near 115 kDa) were present in the cell lysates expressing S(H120), S(H120-S2′/HV80) or S(HV80-H855Q) protein, and no band was detected near 80 kDa (Figure 6E). The protein band intensity was quantitated by Image J software, and the cleavage efficiency was assessed. The result of grey-scale analysis of protein bands showed all proteins were cleaved at S1/S2 cleavage site, the cleavage efficiency at S1/S2 site in S(HV80), S(HV80-R687P) and S(HV80-V692F) groups were significantly higher than that in S(H120) group (Figure 6F). Compared to S(HV80), the cleavage efficiency of one acid amino substitution groups, such as S(HV80-R687P) and S(HV80-V692F), did not cause significant changes (Figure 6G), but the cleavage efficiency of S(HV80-H855Q) showed significant decrease.

**FIG 6.**
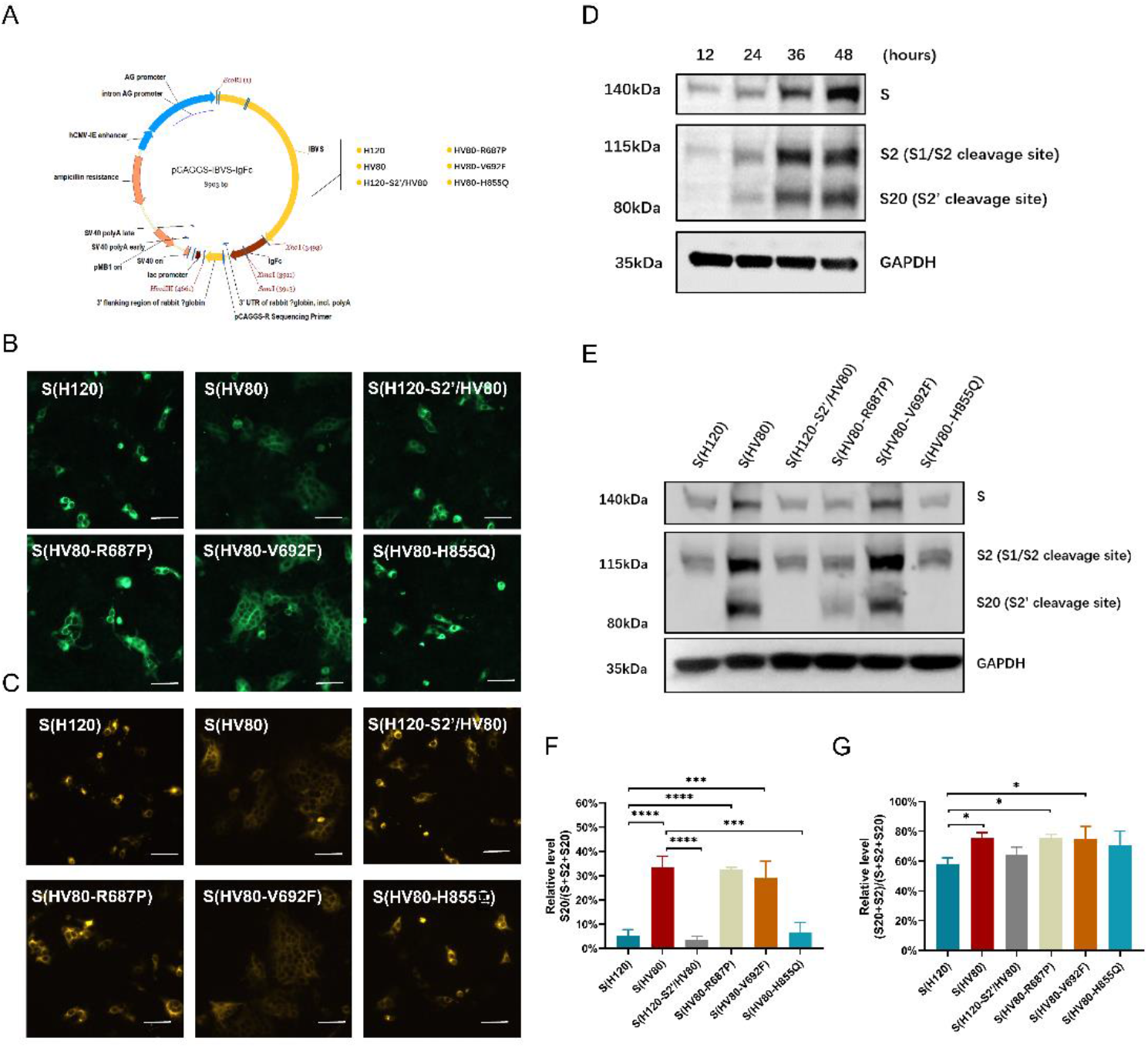
Cell-to-cell fusion induced by different S glycoprotein constructs and proteolytic cleavage of different IBV S glycoproteins. (A) Schematic diagram of recombinant plasmid construction. Upon cleaving the vector pCAGGS-MCS with restriction enzymes (*Eco*RI and *Xho*I), different IBV fragments were inserted and co-expressed with Fc-tag. (B and C) Vero cells were transfected with different S glycoprotein constructs. After 36 h post-transfection, the cells were fixed with cold methanol and permeabilized using 0.5% Triton X-100. The transfected cells were immunolabeled with anti-IBV serum and secondary antibody anti-chicken IgY (IgG) (whole molecule)-FITC antibody produced in rabbit (B) or directly with secondary antibody Goat Anti-Human IgG Fc (DyLight 650) preadsorbed (C). The green (B) or orange (C) fluorescence and the fusion of cells transfected with pCAGGS-IBVS-Fc plasmids were visualized under a fluorescent microscope. (D) Vero cells were transfected with S(HV80) fused with Fc-tag constructs. At 12, 24, 36 and 48 h post-transfection, the cells were lysed and cell lysates were blotted with Goat Anti-Human IgG Fc (HRP) antibodies. The same membrane was also probed with anti-GAPDH monoclonal antibody as a loading control. (E) Vero cells were transfected with different pCAGGS-IBVS-Fc plasmids, respectively. At 36 h post-transfection, cells were harvested and lysates prepared. The viral protein expression was analyzed by Western blot with Goat Anti-Human IgG Fc (HRP) antibodies. The same membrane was also probed with anti-beta-actin monoclonal antibody as a loading control. (F) and (G) The cleavage rate of S1/S2 and S2’ cleavage site. The western blotting protein bands of (E) were analyzed for grayscale values using Image J software, and the cleavage efficiency was assessed and calculated.

## DISCUSSION

This study investigated the mechanism of Vero cell adaptation of the IBV HV80 strain, which is a new adapted strain obtained in previous studies in our laboratory. The target sites that were associated with Vero cell adaptation were performed through different recombinants (chimeric expression of S, S1, S2, or S2′ cleavage site, or other amino acid substitutions) constructed using a reverse genetic system. Virus infection tests and S glycoprotein expression tests showed that the RRRR_690_/S motif at the S2′ cleavage site played a key role in invasion of Vero cells, but PRRR_690_/S at the S2’ cleavage site could be acitivated by proteases in the process of cell-to-cell fusion. Q855H substitution induced the formation of syncytia, and Q at position 855 did not affect the cell invasion of HV80, but hindered the cell-to-cell fusion by blocking activation of the S2’ cleavage site.

S glycoprotein plays an essential role during infection with CoVs *in vitro* and *in vivo*. Exchange of S glycoprotein might result in altered tropism *in vivo* (44). By construct the recombinants, which chimeric expressing a heterologous S glycoprotein, some studies have demonstrated that the cellular tropism of IBV is determined by the S glycoprotein (45, 46). In this study, the chimeric recombinant strain rHV80-S/H120 expressing S glycoprotein from H120 strain lost the ability to infect Vero cells, which demonstrated that S glycoprotein determined the Vero cell tropism of strain HV80. Except for the mutation on S gene, there are a total of 8 amino acid substitutions occurring in replicase 1a gene and E gene. Because it lacked a recombinant, which expressing S glycoprotein of strain HV80 as the genome backbone of strain H120. These results could not rule out an effect of the mutations at other positions on cellular tropism. Two subunits S1 and S2 are generated from S glycoprotein during viral infection. S1 binds to the host receptors, while S2 mediates the fusion of viral and cellular membranes (47). Subsequently, S2 of the Beaudette S glycoprotein was found to be associated with the ability to grow in Vero cells (28). In our previous studies, S2 subunit also determined the primary CK cell tropism of IBV YZ120 strain (46). *In vivo*, exchanging the hypervariable regions on S1 subunit and retaining S2 subunit did not change the tissue tropism of IBVs (48). In this study, we constructed chimeric recombinants expressing the S1 or S2 subunit from the H120 strain, with a genomic backbone of rHV80. The chimeric recombinant expressing the S2 subunit from H120 lost the ability to infect and replicate in Vero cells, which demonstrated that the amino acid changes in S2 were associated with Vero cell tropism of strain HV80. Exchanging the S1 subunit from H120 strain did not affect viral infection and replication in Vero cells. This suggested that the S1 subunit of strain H120 already had the ability to bind to receptors or attachment factors and attach to Vero cell membranes. Therefore, the change in Vero cell tropism of strain HV80 was independent of amino acid substitutions on S1 subunit.

One or two cleavage sites (S1/S2 and S2′) have been found in the S glycoprotein of CoVs, which are cleaved by an appropriate protease on the plasma or endosomal membrane. The S1/S2 cleavage site is located at the S1 and S2 interface, and the tertiary structure indicates that it is exposed on the dorsolateral surface of the S glycoprotein. The S2′ cleavage site is restrained on the inside of trimeric S glycoprotein by subunit S1, and the cleavage of this site is triggered upstream of the FP (38) (Figure 7A and B). In SARS-CoV, the furin cleavage of S glycoprotein enhances cell-cell fusion but does not affect virus entry (49). Two furin cleavage sites (S1/S2 and S2′) have been identified on the S glycoprotein of MERS-CoV. Cleavage at S1/S2 occurs during S biosynthesis in Vero producer cells, and S2′ cleavage occurs during virus entry into target cells (26). Unlike other IBVs, the Beaudette strain possesses two furin cleavage sites, while other IBVs only possess the S1/S2 cleavage site. rIBV with mutation or deletion at the S1/S2 furin site also infects Vero cells but forms smaller syncytia than the wild-type virus. This suggests that furin cleavage at the S1/S2 site is not necessary for, but can promote, infectivity and syncytial formation of IBV in Vero cells (50). No syncytia in the cells transfected with S glycoprotein with mutation or deletion at the S2′ cleavage site. They also demonstrated that the S2′ cleavage site resulted in the expansion of cell tropism, and XXXR_690_/S motif was likely the minimal sequence required to support IBV infectivity in Vero cells (50). In this study, this cleavage site (RRRR_690_/S) also appeared on the S glycoprotein of strain HV80 with the substitution P687R. Furin and general proprotein convertases (PCs) activitiy of the motifs at the S2′ cleavage site (from position 684 to 692) were predicted on the ProP-1.0 server (https://services.healthtech.dtu.dk/service.php?ProP-1.0). It determined that both the furin and general PCs predict scores of the motif PSSRRRR_690_/SF at S2’ cleavage site were higher than that of the motif PSSPRRR_690_/SF (Table 2). The rHV80-S2′/H120 strain, with the substitution of R687P in S glycoprotein (motif: PRRR_690_/S) lost the ability to infect Vero cells. Bearing a furin-recognition site at S2′ cleavage site on S the glycoprotein of strain H120 conferred upon the H120-S2′/HV80 strain the ability to infect Vero cells, but the replication and transmission efficiency were limited. These results demonstrated that the S2′ site (RRRR_690_/S) might trigger the adaptation of HV80 to Vero cells by furin. No plaques appeared in Vero cells incubated with the rHV80-S(R687P) strain because no infection occurred. Alternatively, some CoVs have acquired mutations in the S glycoprotein with the multi-basic motifs of XBBXBX or XBXXBBBX (B: a basic residue) during serial passage in cells, enabling novel binding capacity to heparan sulfate (HS) at the cell surface (51, 52). The Beaudette strain harbors an HS binding motif _685_SSRRRRSV_692_ at the S2′ cleavage site of S glycoprotein, which reveals that HS binding to the S glycoprotein allows the Beaudette strain to attach to cell surfaces (21). Amino acid sequence of _685_SSRRRRSV_692_ at the S2′ site of strain HV80 is accordance with XBBXBX of the HS binding motif. Therefore, we speculated that the expansion of Vero cell tropism also might be associated with formation of the HS binding motif at the S2′ cleavage site. Either due to furin cleaving or HS binding, only possess RRRR_690_/S motif at S2’ cleavage site on S glycoprotein did not lead to high efficiency of infection and replication of the rH120-S2′/HV80 strain in Vero cells. Other mutation sites on the S2 subunit or the genome backbone might promote viral infection and replication in Vero cells.

**FIG 7.**
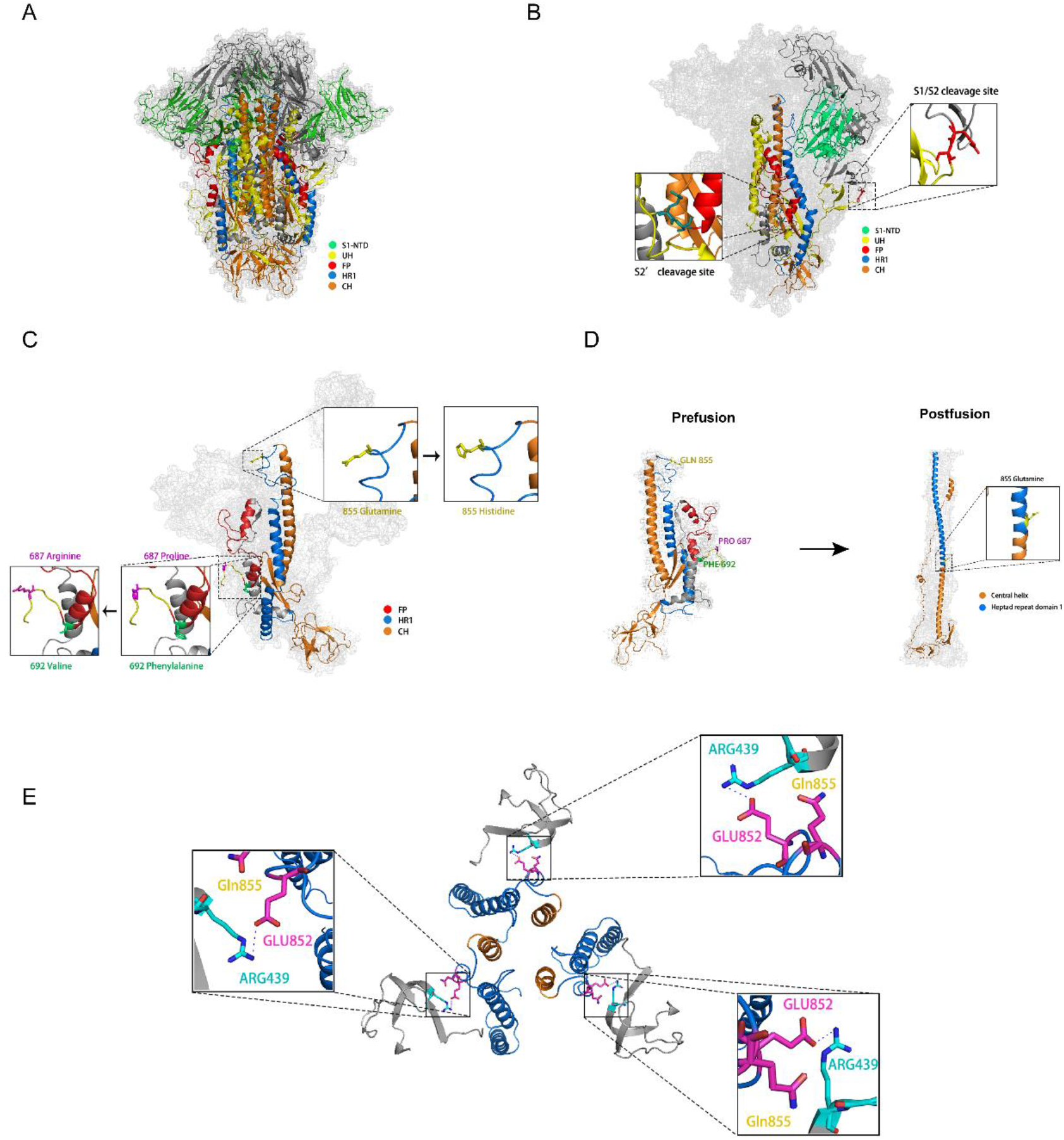
Structure and schematic of the IBV S glycoprotein. (A) IBV S glycoprotein trimeric structure modelled in PyMol predicted by the Swiss-model referred to 6cv0.1. S1-NTD, L, fusion peptide (FP), heptad repeat domain 1(HR1) and central helix (CH) are shown in light green, yellow, red, blue and orange, respectively. (B) The monomer structure of the S glycoprotein in the prefusion state. The S1/S2 and S2′ cleavage sites are presented within the black squares. (C) S2 domain in a prefusion form. The mutations of P687R, F692V and Q855H are labeled as sticks in magenta, green and yellow, respectively, and magnified within the black squares. (D) S2 domain from a prefusion form to postfusion form. The postfusion forms were predicted by the Swiss-model referred to 6xRA. The amino acids at position 855 in postfusion forms were labeled as sticks and magnified within the black squares. (E) The structure of salt-bridge in Glu852-ARG439. The dotted lines represent salt-bridge.

**TABLE 1.** Names of recombinant viruses corresponding to the names of plasmids pMDVM8.

| Viruses | Plasmid pMDVM8 |
| --- | --- |
| rH120 | pMDVM8-H120S |
| rHV80 | pMDVM8-HV80S |
| rHV80-S1/H120 | pMDVM8-H120S1HV80S2 |
| rHV80-S2/H120 | pMDVM8-HV80S1H120S2 |
| rH120-S2'/HV80 | pMDVM8-H120SHV80S2' |
| rHV80-S2'/H120 | pMDVM8-HV80SH120S2' |
| rHV80-S(V692F) | pMDVM8-HV80S(V692F) |
| rHV80-S(N802K) | pMDVM8-HV80S(N802QK) |
| rHV80-S(R822Q) | pMDVM8-HV80S(R822Q) |
| rHV80-S(H855Q) | pMDVM8-HV80S(H855Q) |
| rHV80-S(L1008V) | pMDVM8-HV80S(L1008V) |
| rHV80-S(F1010Y) | pMDVM8-HV80S(F1010Y) |

**TABLE 2.**
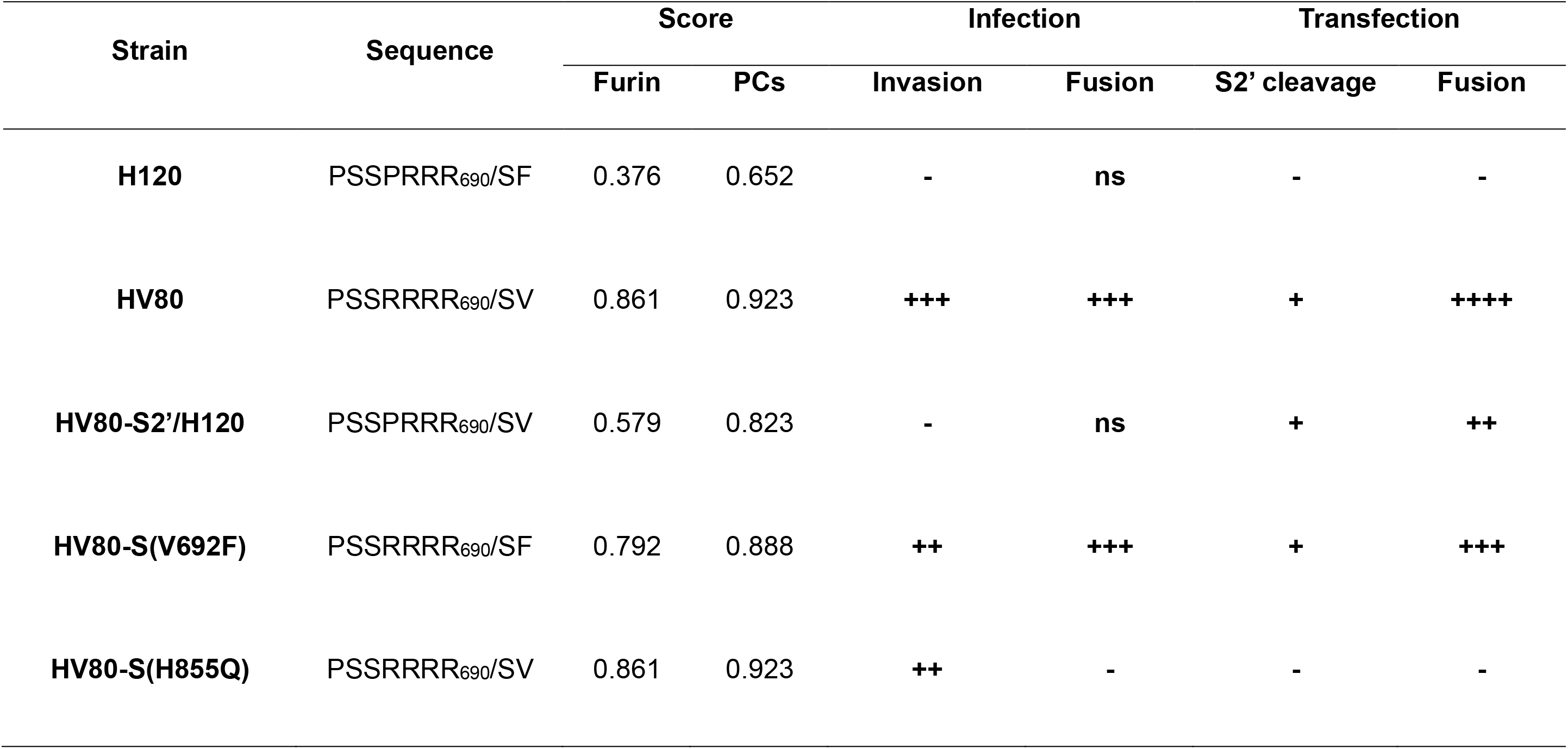
Predicted cleavage by furin and general PCs at the S2′ cleavage site of different strains and the cleavage, infection and fusion in Vero cells infected with different strains.

As a class I viral fusion protein, it mediates membrane fusion triggered by several factors, such as receptor binding, proteolysis of S glycoprotein, or low pH. Subsequent to proteolysis, exposed FP inserts into the cell membrane. Trimeric HR2 folds back to trimeric HR1, envelops the HR1, and together they form a six-helix bundle. This promotes conformational rearrangement of the viral and cell membranes, and brings them into fusion (36, 37, 53). The results of our sequence analysis showed that F692V substitution was located on the FP; K802N, Q822R and Q855H substitutions were located on HR1; and V1008L and Y1010F were located on the central helix (CH) domain (Figure 1). Substitution of each amino acid site affected viral replication in Vero cells, and replication efficiency in the V692F group showed the most obvious decline. F692V substitution was adjacent to site 687, but the latter formed an S2′ furin cleavage site, and the former was located at the apex of the putative FP domain (Figure 7C). The plaque size of Vero cells infected with the rHV80-S(V692F) strain was not affected, but the number of plaques per well formed by this strain in Vero cells was lower than that in other groups infected with the same viral copy number. Although the viral RNA copy number in Vero cells infected with the rHV80-S(V692F) strain was lower at the early stage of infection, large syncytia were seen in the infected cells. Prediction of protease activity at the S2′ cleavage site showed that F692V substitution enhanced furin proteolysis activity at the S2′ site (Table 2). It indicated that viral invasion of the host cells but not cell-to-cell fusion might be influenced by the V692F substitution through an effect on proteolysis at the S2′ cleavage site. Notably, substitution H855Q resulted in a loss of cell fusion for viruses, and no syncytia was appeared in Vero cells infected with rHV80-S(H855Q). The viral growth curve for rHV80-S(H855Q)-infected cells was different from that of other groups. The viral copy number in the culture supernatant of rHV80-S(H855Q)-infected cells was higher in early infection, but the slope for proliferation was less than the growth curve of other strains. We speculated that substitution of H855Q resulted in loss of ability to induce cell-to-cell fusion in infected Vero cells. The viral products from infected cells could not spread to uninfected neighboring cells by cell-to-cell fusion. In some single cells infected with the rHV80-S(H855Q) strain, newly synthesized virions were released into the culture supernatant. These viruses bound to the surface of other uninfected cells, and initiated a new round of infection. Although viral RNA copies of rHV80-S(H855Q) could be detected in cell culture supernatant in early infection using this transmission mode, the total efficiency of transmission and infection were lower than those for viral transmission through cell-to-cell fusion. The plaque size in the rHV80-S(H855Q)-infected group was significantly smaller than in the other infected groups. For CoVs, mutations in the HR1 region have been linked to changes in viral tropism or are associated with its adaptive evolution (40, 54). Leucine to phenylalanine substitution at position 857 from p65 strain of Vero-adapted IBV converted the nonfusogenic S glycoprotein to fusogenic (42). Interestingly, the positions of L857F substitution of Beaudette p65 strain is closed to that of H855Q substitution of HV80 strain, both of them are located on the C-terminal end of HR1.

Proteolysis of CoV spikes can lead directly to membrane fusion and thereby serves as an essential trigger for membrane fusion (4, 55). To further understand whether the substitution at the S2′ cleavage site could have effects on cell-to-cell fusion, we constructed S glycoprotein eukaryotic expression plasmids fused with an EFGP or Fc tag. Large syncytia appeared in Vero cells expressing S(HV80), S(HV80-N802K), S(HV80-R822Q), S(HV80-L1008V) or S(HV80-F1010Y) fused with an EGFP tag. However, only individual cells with high fluorescence intensity were present in Vero cells expressing S(H120-S2’/HV80) or S(HV80-H855Q) fused with an EGFP tag, which confirmed that Q855H substitution improved the efficiency of viral infection by promoting cell-to-cell fusion. These results are consistent with the aforementioned results of syncytia formation in recombinant-infected Vero cells. Both IBV rH120 and rHV80-S(HV80-R687P) could not infect Vero cells. No cell-to-cell fusion occurred in Vero cells transfected with pCAGGS-S(H120)-EGFP, while the cells transfected with pCAGGS-S(HV80-R687P)-EGFP were present as individual cells and some small syncytia with high fluorescence intensity. This indicated that the motif PRRR_690_/S at the S2′ cleavage site could still induce cell-to-cell fusion in transfected cells. Less cell-to-cell fusion occurred in cells expressing S(HV80-V692F) glycoprotein fused with EGFP-tag. However, large syncytia were observed in Vero cells transfected with pCAGGS-S(HV80-V692F)-Fc. The culture supernatant was harvested to extract viral RNA. After being amplified and sequenced, no other mutation, except that encoding V692F, occurred on S gene. We speculated that the EGFP tag might have hindered the fusion function of S(HV80-V692F) glycoprotein. This indicated that the ability of rHV80-S(V692F) to invade Vero cells was reduced, but cell-to-cell fusion induced by this strain was not affected by V692F substitution.

To gain further insight into S glycoprotein cleavage of different recombinants, which transient expressing and fusing in Vero cells, cell lysates were collected separately after transfection. Western blotting showed that, all proteins were cleaved at S1/S2 cleavage site, the S glycoprotein which can induce cell-to-cell fusion, such as S(HV80), S(HV80-R687P) and S(HV80-V692F), showed higher cleavage activity at S1/S2 cleavage site. Unlike S(H120) protein, S1/S2 and S2′ cleavage sites of S(HV80) protein were cleaved into two S2 subunit fragments of different sizes that were expressed in Vero cells and induced cell-to-cell fusion. Whether P to R substitution occurred at position 687 of the S2′ cleavage site had little effect on cleavage at the S2′ site. Only possess of RRRR_690_/S motif, the S(H120-S2’/HV80) glycoprotein could not be activated at the S2′ cleavage site. Although weaker protein bands were detected, single substitution of R687P on the S(HV80-R687P) glycoprotein still activated the S2′ cleavage site. The efficiency of cleavage at the S2’ cleavage site between the S(HV80-R687P) and HV80 groups did not show a significant difference. Nonetheless, the exact mechanism requires further investigation. But compared to S(HV80) and S(HV80-V692F), weaker bands of S glycoprotein were detected in S(HV80-R687P) expression group. In IFA, the transfected positive cells were present as individual cells and some small syncytia with high fluorescence intensity, and some of them might have disintegrated, denatured and dropped off. The weaker bands of S glycoprotein also detected in cells expressing S(H120), S(H120-S2’/HV80) and S(HV80-H855Q), which also failed to induce cell-to-cell fusion. We speculated that the weaker bands detected in cells expressing S(HV80-R687P) were due to disintegration of positive cells at late phases of transfection.

Proteolysis of the S2′ cleavage site was affected by H855Q substitution. The S2′ cleavage sites of all S glycoproteins with histidine at position 855 were activated, while no cleavage occurred at S2′ sites with glutamine at the same position. As a result, we speculate that there are two possible explanations for this phenomenon. The first is the motif of the S2’ cleavage site, rather than the cleavage event, is associated with syncytium formation. It is likely that the exposed S2’ cleavage site played a role in cell-to-cell fusion through other means. Another explanation is that cell-to-cell fusion event is triggered by the cleavage at S2’ cleavage site of extremely low amounts of S glycoprotein, but whether fusion occurs is also determined by the conformation change, which is affected by the amino acid substitution at position 855.

The above results were collated into a table to analyze the correlation between amino acid substitution at positions 687, 692 and 855 and viral infection and cell fusion. It showed that in the process of viral infection, the ability to invade Vero cells was consistent with the furin scores of the motif at the S2′ cleavage site of S glycoprotein. The higher the furin activation at the S2′ cleavage site, the stronger the ability to invade Vero cells. The scores of motif PSSPRRR/SX (X: F or V) were lower, and P687R substitution directly blocked invasion of Vero cells by IBV. However, the S2′ cleavage site with PRRR_690_/S motif was still cleaved, and its S glycoprotein could still induce cell-to-cell fusion in the transfected Vero cells. It might be mediated by other proteases (e.g., PCs), but the detailed mechanism remains to be established. The substitution of F692V enhanced the proteolytic activity of furin at the S2′ cleavage site. The ability of V692F substitution recombinants to invade Vero cells decreased with virus-to-cell fusion. In the process of viral infection, virus-to-cell fusion was not affected by the substitution at position 855, but was activated by the motif RRRR_690_/S at the S2′ cleavage site. However, there are no data to show that the S2′ cleavage site was cleaved in the process of HV80 infection. In transient transfection assays, we observed that cell-to-cell fusion occurred through proteolysis at the S2’ cleavage site. However, this process was blocked by the H855Q substitution. The tertiary structure of the S glycoprotein showed that 855 site was located outside the HR1 helix, between the HR1 and CH helices. In the prefusion state, constrained by the S1 subunit trimer, the amino acid at position 855 was present in the form of loops, and it made the HR1 and CH helices fold in the center of the S2 trimer. Upon fusion triggering, conformation of the S2 subunit changed, with “jack-knife” refolding of the HR1 helix and intervening regions into a single continuous helix appended to the CH (Figure 7D). Studies on the structure and function of the SARS-CoV-2 S glycoprotein have revealed the presence of various intermolecular forces in its monomers, including salt bridges and hydrogen bonds. Of these, the salt bridge between D614 and K854 on the S glycoprotein is the most well-understood. This bridge is critical for triggering conformational changes following receptor binding. During the early stages of the SARS-CoV-2 epidemic, the D614G substitution caused a disruption in the salt-bridge interaction with K854, resulting in increased transmission efficiency of new variants(56). Structural studies on the S glycoprotein of the alpha variant revealed that the mutations A570D and S982A could impact RBD conformational changes. This is due to the addition of a K852-D570 salt bridge and loss of stabilizing interactions afforded by the hydrogen bonds between S982 and G545, T547(57). Previous studies have shown that the L857F substitution on the S glycoprotein of the IBV Beaudette strain is linked to virus-induced syncytium formation, while the E405D substitution has a compensatory effect. However, the lack of crystal structures for the IBV S glycoprotein has made it unclear whether these two sites are directly related to each other. The H120 S glycoprotein structure was generated through homology modeling utilizing the Swiss-Model server (swissmodel.expasy.org), while the VMD software was employed to predict the salt bridges. The model reveals a salt bridge interaction between E852 and R439, and the site at position 855 was adjacent to this salt bridge. We speculate that the stability of the S glycoprotein is linked to the salt-bridge interaction. Additionally, the Q855H substitution may affect the stability of the E852-R439 salt bridge, leading to more easily triggered conformational changes in the S glycoprotein. After the conformation changing, the S2′ cleavage site is exposed for activation by proteases. In other groups with no H855Q substitution, the cell-to-cell fusion efficiency was dependent on the motif at the S2′ cleavage site. We found that there was a different mechanism between virus-to-cell fusion and cell-to-cell fusion mediated by HV80 S glycoprotein, which needs to be further studied.

A phenomenon described as receptor-independent spread was shown by the mouse hepatitis virus (MHV)-JHM strain. The MHV hepatotropic strains infect host cells by binding to the CEACAM1 receptor (58), but MHV-JHM strain can spread from infected mouse cells to cells lacking mCEACAM1a, and it can infect and kill CEACAM1a^−/−^ mice (59). It has been speculated that the MHV-JHM strain can potentially use an alternative, less-effective receptor to initiate infection (5). Once the virus had triggered a primary infection in mouse nerve cells, the JHM strain can spread rapidly by cell-to-cell fusion rather than binding to receptors on the surface of the cell membrane (60). We do not think that there is a natural protein receptor on Vero cells, and the HV80 strain may use other less-effective receptors or attachment factors (e.g, sialic acid) to invade Vero cells. The virus acquired a more effective S2′ cleavage site motif or HS binding site by P687R substitution, and successfully infected the Vero cells by inducing virus-to-cell fusion. The F692V substitution further enhanced the proteolytic activity of furin at the S2′ cleavage site. The Q855H substitution occurred in the process of viral adaptation to Vero cells, which conferred upon S glycoprotein expressed on the cell surface the ability to fuse with neighboring cells. Once primary infection is established in Vero cells, the HV80 strain can trigger more efficient infection through rapidly forming large syncytia by its S2 subunit with high fusion capacity. The detail mechanisms need to be further studied.

In conclusion, the substitution of P687R and F692V in S glycoprotein formed a highly efficient S2’ cleavage site of HV80 strain, which enabled the virus to invade Vero cells. The mechanisms were different between virus-to-cell and cell-to-cell fusion induced by HV80 S glycoprotein. In the process of HV80 virus infection, the virus-to-cell fusion in Vero cells were caused by P687R substitution, but cell-to-cell fusion was associated with Q855H substitution in S glycoprotein. Whether the S2’ site cleaves even depends on the substitution of Q855H in cell-to-cell fusion induced by S glycoprotein. Finally, a number of important limitations need to be considered. First, the effects on viral adaption to Vero cells by other mutations in replicase gene 1a and E could not be excluded. Second, it is unclear whether S2’ site be cleaved during the invasion of the HV80 strain into Vero cells. Third, it is unclear how S2’ cleavage site and Furin exert their action in cell-to-cell fusion induced by S glycoprotein. This study provides new theoretical insights into mechanisms of IBV adaptation in mammalian cell lines.

## MATERIALS AND METHODS

### Viruses and cells

Vero cells were purchased from *Shanghai Cell Bank, Chinese Academy of Sciences (Shanghai, China)*, and cultured in Dulbecco’s modified Eagle’s medium (DMEM, *Gibco, Shanghai, China*) supplemented with 10% inactivated fetal bovine serum (FBS, *Gibco*) and 1% penicillin–streptomycin (*Beyotime*). All specific-pathogen-free (SPF) embryonated eggs were purchased from Beijing Boehringer Ingelheim Vital Biotechnology. Viruses were propagated in 10–11-day-old SPF embryonated eggs. (i) HV80, a Vero-cell-adapted strain, which was serially passaged for five generations in chicken embryos, 20 passages in CK cells and 80 passages in Vero cells(43). (ii) rHV80, a molecular clone of the HV80 strain. (iii) rH120, a molecular clone of the widely used vaccine H120 strain. (iv) rHV80-S/H120, rHV80-S1/H120 or rHV80-S2/H120, expressing the chimeric S glycoprotein from the H120 strain, or the chimeric S glycoprotein composed of the S1 subunit derived from the H120 strain and S2 subunit from the HV80 strain, or the S1 subunit derived from the HV80 strain and S2 subunit from the H120 strain, with the backbone genome of the HV80 strain. (v) rH120-S2′/HV80 or rHV80-S2′/H120, expressing or knocking down the S2′ cleavage site, with the genome of the rH120 or rHV80 strains. (vi) rHV80-S(V692F), rHV80-S(N802K), rHV80-S(R822Q), rHV80-S(H855Q), rHV80-S(L1008V) and rHV80-S(F1010Y) were based on the backbone genome of the rHV80 strain, with nucleotides on the S gene representing H120 amino acid substitutions V692F, N802K, R822Q, H855Q, L1008V and F1010Y, respectively.

### Construction of recombinant viruses

Construction of different recombinant viruses is described in the schematic illustration in Figure 1. The full-length cDNA clones of IBVs were constructed as described previously (61)(62). According to the distribution of IIs restriction enzymes *Bsa*I on the genome of the HV80 strain, fragments VM1–VM10 of HV80 genome were amplified by PCR. The sequences of primers have been described previously (61). cDNA of nucleoprotein gene was amplified using plasmid pVAXN as a template and the following primer pair: NT7S (5’-CCACTGCTTACTGGCTTATCG-3’) and NT7A (5’-TTTTTTTTTTTTTTTTTTTTTTTTTAGGAAAGGA CAG-3’). The PCR products were ligated into pMD19-T vector, transformed into *Escherichia coli* to obtain plasmids pMDVM1–pMDVM10. Fragments VM1–VM10 were recovered from the recombinant plasmid ligated by *Bsa*I. Full-length genomic cDNA was obtained by ligating the 10 fragments *in vitro*. After transcribing using the In vitro Transcription T7 Kit (TaKaRa), BHK-21 cells were cotransfected with both full-length genomic and nucleoprotein gene RNA. The cell culture was collected 48 h after transfection, freeze-thawed three times, and inoculated into 10-day-old SPF chicken embryos. The rescued molecular clone strain rHV80 was obtained from the allantoic fluid at 48 h post-inoculation. Both strains with the H120 genome backbone and HV80 backbone shared one set of primers and were rescued according the same method. For other recombinants, the difference was the construction of different plasmids pMDVM8, which need to modified and amplified the fragment VM8 (S gene) (Table 1). In the In-Fusion PCR cloning system (Clontech), the S1 or S2 gene of the HV80 strain was replaced with the corresponding region of the H120 strain to construct the recombinant plasmids pMDVM8-HV80S1H120S2 and pMDVM8-H120S1HV80S2, which contained the chimeric S genes. pMDVM8 plasmids for the recombinants harbored one-point amino acid substitution in the S gene of the HV80 strain and were amplified and constructed using overlapping PCR.

### Indirect immunofluorescence staining

A monolayer of 2 × 10^5^ Vero cells per well was seeded in six-well plates and cultured for 18 h. The cells were washed three times with PBS and incubated with different viruses at 10^7^ viral RNA copies at 37℃ and 5% CO_2_. After 1-h incubation, viral inoculum was removed and cell monolayers were washed with PBS three times to eliminate unbound virus, and then cultured in DMEM containing 1% FBS. After 24- and 48-h incubation, the infected cells were fixed in cold methanol for 30 min and 5% Triton-X-100 in PBS for 20 min. The treated cells were blocked with 5% nonfat dry milk for 2 h, and incubated with anti-IBV serum as the primary antibody (1:100) at 4℃ overnight. After washing with PBS three times, the treated cells were performed using anti-chicken IgY (IgG) (whole molecule)-FITC antibody produced in rabbit (Sigma–Aldrich, dilution 1:640) as the secondary antibody. Cells were further washed in same way, and the nuclei were stained with 4′6-diamidino-2-phenylindole (DAPI) (Beyotime Biotechnology). Images were taken using a Leica DMi8 microscope (Leica, Germany).

### Assessment of viral growth curve

Vero cells were seeded at 2 × 10^5^ cells/well in 12-well plates 18 h prior to infection with rIBVs. Cells were washed three times with PBS and infected with different viruses at 5×10^5^ viral RNA copies per well on the following day. Three replicate wells were included for each virus. After 1-h incubation, the infected cells were washed three times with PBS again, and cultured in 1 ml/well DMEM containing 1% FBS. Cell culture supernatants (70 μl per well; total 210 μl per group) of were collected at 12, 24, 36, 48, 60 and 72 hpi and stored at −70℃. Viral replication in Vero cells was assessed by quantitative determination of viral RNA as follows. Viral RNA was extracted from the supernatant using a Virus DNA/RNA Extraction Kit 2.0 (Vazyme), and reverse transcribed into cDNA using PrimeScript RT Master Mix (Perfect Real time) (TaKaRa). cDNA copies of IBVs were quantified by real-time PCR using TB Green Premix Ex Taq II (Tli RNaseH Plus) (TaKaRa) and the following primer pair: DL2S, CCGTTGCTTGGGCTACCTAGT; DL2A, CGCCTACCGCTAGATGAACC. The viral RNA copies were calculated by generating a standard curve using serial dilutions of DNA sequence for the 1a gene.

### Eukaryotic expression plasmid construction

To analyze the activation for cell–cell fusion of different S glycoproteins, pCAGGS-IBVS-EFGP/Fc eukaryotic expression plasmids, which encoded full-length S gene from different viruses, were constructed. The fragments EGFP or Fc were amplified and ligated to the pCAGGS-MCS vector by different restriction enzymes (EGFP: *Sac*I and *Xho*I, Fc: *Xho*I and *Bsg*I) to construct the plasmids pCAGGS-EGFP or pCAGGS-Fc. PrimeSTAR polymerase (TaKaRa) was used to amplify the S gene sequence by PCR with the following two sets of primers: forward, CCGGAATTCATGTTGGTAACACCTCTTTT, and reverse: GGCGAGCTCAACAGACTTTTTAGGTCTGT (IBVs fused with EGFP), forward, CCGGAATTCATGTTGGTAACACCTCTTTT, and reverse, GGCCTCGAGAACAGACTTTTTAGGTCTGT (IBVS fused with Fc). PCR products were purified by agarose gel electrophoresis, and digested with two sets of restriction enzymes (EGFP: *Eco*RI and *Sac*I, Fc: *Eco*RI and *Xho*I). T4 ligase (TaKaRa) was used to connect the digested PCR products and the vector at 16°C for 2 h. Ligase products were transformed into JM109 cells. After transformation, monoclonal cell strains were selected for sequencing. Positive plasmid extracted from clones with correct sequence using TIANprep Mini Plasmid Kit II (Tiangen).

### Cell transfection assay

Vero cells were seeded in 35-mm dish 18 h prior to transfection, and grown to 90% confluence on the next day. Vero cells were transfected with pCAGGS-IBVS-EGFP/Fc using Lipo3000 (Invitrogen) and Opti-MEM (GIBCO). Plasmid DNA (5 μg) was mixed with 7.5 μl Lipo3000 and diluted in 250 μl Opti-MEM before addition to cells. After 15-min incubation at room temperature, the complex was added to the cells with complete DMEM containing 10% FBS and incubated at 5% CO_2_ and 37℃. At 36 h post-transfection, the green fluorescence and fusion of cells transfected with pCAGGS-IBVS-EGFP plasmids were visualized under a fluorescence microscope (Leica DMi8, Germany)). Vero cells transfected with pCAGGS-IBVS-Fc plasmids were lysed and total protein was harvested and estimated for western blotting after 36 h transfection.

### Western blotting

Cells were lysed using RIPA Lysis Buffer (Beyotime) containing 1 mM phenylmethylsulfonyl fluoride protease inhibitor. The samples were incubated at 4℃, and collected for centrifugation at 13 000 rpm for 10 min. SDS loading buffer (Beyotime) was added and the samples were boiled for 10 min. The samples were run on precast SurePAGE gels (Bis–Tris, 10×8, 4%– 12%, 15 wells; GenScript) and transferred to polyvinylidene difluoride membranes (Beyotime Biotechnology) using the Bio-Rad Transblot transfer system. The membranes were blocked with 5% nonfat dry milk *overnight* at 4℃, and directly incubated with the secondary antibody using Goat Anti-Human IgG Fc (HRP) preadsorbed (Abcam). After washing five times with Tris-buffered saline and Tween 20, the membranes were exposed to Super Signal ECL (Pierce) and the chemiluminesence signal was captured using a ChemiDoc Imagining system (BioRad). The western blotting protein bands were analyzed for grayscale values using Image J software, and the cleavage efficiency was assessed. It was calculated using the following formula: S1/S2 cleavage rate = (S2 gray value + S20 gray value) / (S gray value + S2 gray value + S20 gray value), S2’ cleavage rate = S20 gray value / (S gray value + S2 gray value + S20 gray value).

### Statistical analysis

GraphPad Prism 7 software (*GraphPad Software Inc., CA, USA*) was used for all data analysis. Experimental data were expressed as mean ± standard deviation, and were analyzed with two-way analysis of variance (ANOVA). P values: \**P* < 0.05, \*\**P* < 0.01, \*\*\**P* < 0.001, \*\*\*\**P* < 0.0001.

## ACKNOWLEDGMENTS

This study was supported by the Natural Science Foundation of Jiangsu Province (BK20201483), and National Natural Science Foundation of China (31602091, 31572524).

